# Neurogenetic identification of mosquito sensory neurons

**DOI:** 10.1101/2022.11.22.517370

**Authors:** Joanna K. Konopka, Darya Task, Danny Poinapen, Christopher J. Potter

## Abstract

*Anopheles* mosquitoes, as vectors for the malaria parasite, are a global threat to human health. To find and bite a human, they utilize neurons within their sensory appendages. However, the identity and quantification of sensory appendage neurons are lacking. Here we use a neurogenetic approach to label all neurons in *Anopheles coluzzii* mosquitoes. We utilize the Homology Assisted CRISPR Knock-in (HACK) approach to generate a *T2A-QF2^w^* knock-in of the synaptic gene *bruchpilot*. We use a membrane-targeted GFP reporter to visualize the neurons in the brain and to quantify neurons in all major chemosensory appendages (antenna, maxillary palp, labella, tarsi). By comparing labeling of brp>GFP and Orco>GFP mosquitoes, we predict the extent of neurons expressing Ionotropic Receptors or other chemosensory receptors. This work introduces a valuable genetic tool for the functional analysis of *Anopheles* mosquito neurobiology and initiates characterization of the sensory neurons that guide mosquito behavior.

**Graphical Abstract:** 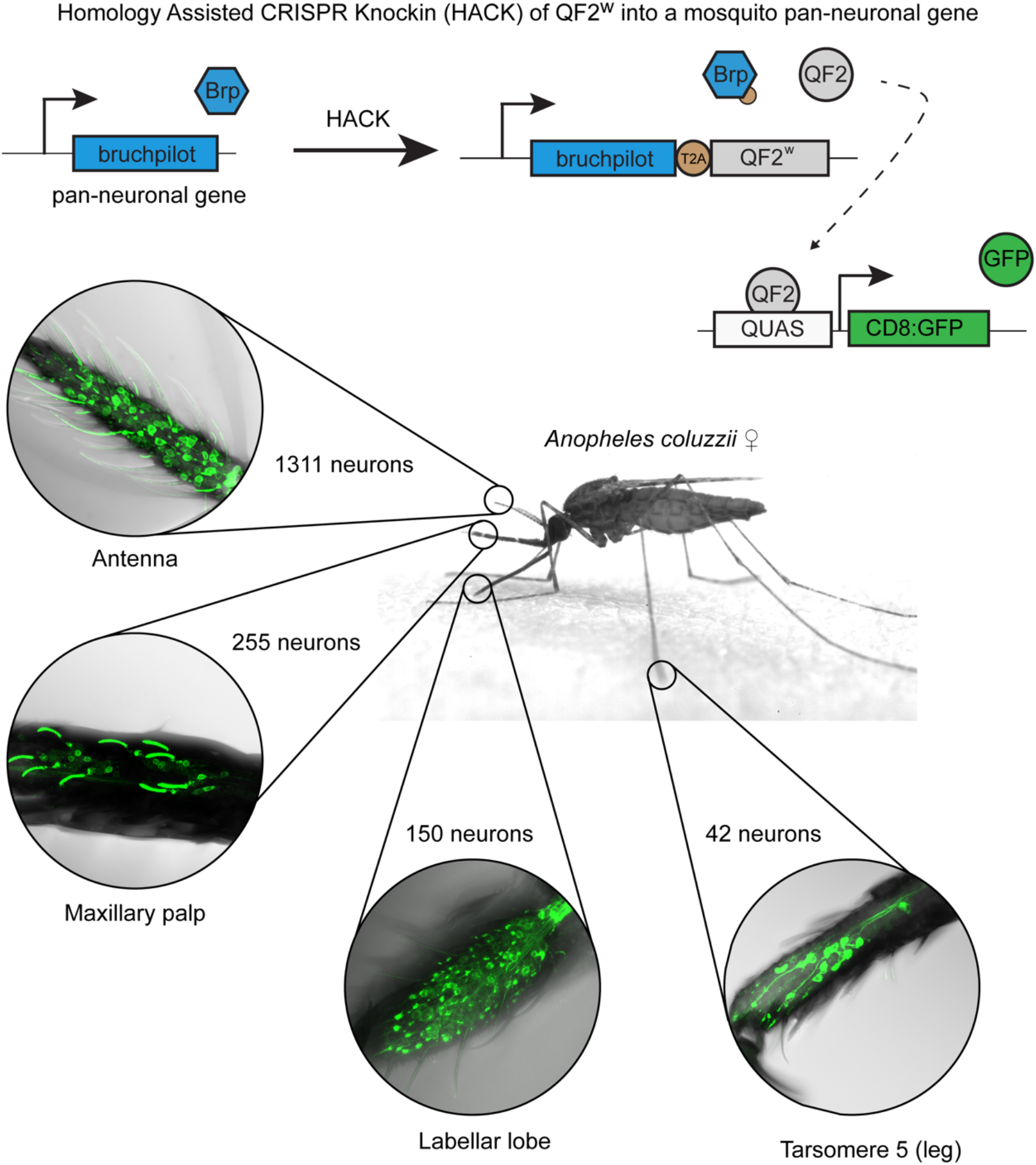

## Introduction

Female mosquitoes are a threat to global human health as they can transmit various diseases through their bites^1–3^. The *Anopheles* species of mosquitoes are the deadliest animals on earth as many prefer to bite humans and can serve as vectors of malaria, which kills ~ 600,000 people each year^4^. Female anthropophilic mosquitoes rely heavily on their finely-tuned sensory systems to forage, mate, locate human hosts, and select oviposition sites^5^. These major behavioral decisions affecting their survival and fitness utilize mechanical (including sound reception), visual, and olfactory cues^5–10^. Olfaction is one of the most critical sensory modalities, involved in many mosquito behaviors, and plays a major role in host seeking^9–12^.

Mosquito behaviors occur in response to stimuli being detected by neurons in sensory appendages, and this information being transmitted, processed, and integrated in the brain. The main mosquito chemosensory appendages involved in host searching and bite site location include antennae, maxillary palps, and the labella of the mouthpart (**Fig. 1A and 1B**). Sensilla (specialized hairs) on the surface of these sensory appendages house neurons. Dendrites of those neurons express receptors that interact with chemosensory cues in the environment. Various combinations of these chemosensory receptors allow for detection and discrimination of a vast number of odorants and other environmental cues, ultimately guiding females in their behaviors. The largest gene families of receptors involved in olfaction are Odorant Receptors (ORs), Ionotropic Receptors (IRs), and Gustatory Receptors (GRs). Current research focuses on the functional analysis of these receptors and creation of mutant mosquitoes targeting individual subunits or chemosensory co-receptors in order to determine their role in mosquito host-searching behavior^13–23^.

**Figure 1.**
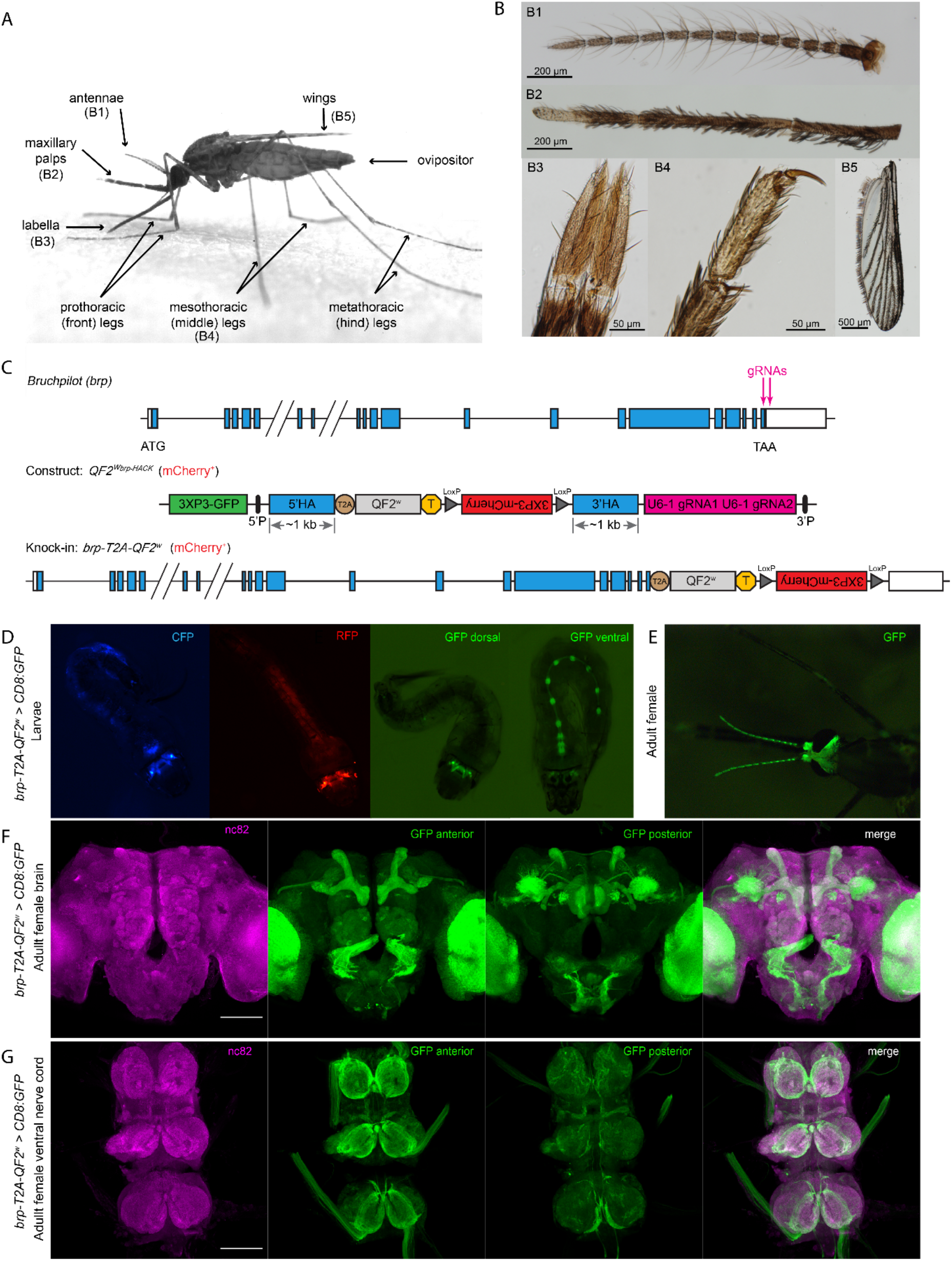
Generation and validation of *brp-T2A-QF2^w^* HACK knock-in driver line for pan-neuronal expression in *Anopheles coluzzii* mosquitoes. (**A**) *An. coluzzii* female mosquito with chemosensory appendages highlighted. (**B**) Chemosensory appendages of the head and the body. (**C**) Schematic diagram of HACK knock-in strategy. Top: *An. coluzzii bruchpilot* (*brp*) target gene with exons indicated as blue boxes and introns as horizontal lines. Diagonal lines indicate truncation of long introns for visualization purposes. Guide RNAs (gRNAs) targeting the stop codon of *brp* gene are indicated by pink vertical arrows. Middle: *QF2w^brp-HACK^* construct with gRNAs (pink), *T2A-QF2^w^* (tan and grey, respectively), floxed *3XP3-mCherry* eye marker (red), and transcriptional stop (T; yellow). Additional *3XP3-GFP* (green) was added upstream of the 5’ homology arm (5’HA; blue) to allow screening against random insertions of the construct in the genome. Bottom: The final knock-in *brp-T2A-QF2^w^* with the construct integrated at the end of *brp* gene. The knock-in can be crossed to a reporter (*QUAS-CD8:GFP*) to examine the endogenous expression pattern of the *brp* gene (**D**) *brp-T2A-QF2^w^* > *QUAS-CD8:GFP* larvae viewed under CFP (blue eye marker for *QUAS-CD8:GFP*), RFP (red eye marker *for brp-T2A-QF2^w^*), and GFP filters. GFP fluorescence indicates expression of *QUAS-CD8:GFP* in the brain and ventral nerve cord in the pattern of *brp* gene. (**E**) *brp-T2A-QF2^w^* > *QUAS-CD8:GFP* adult female with GFP fluorescence in the antennae and brain (visible through the cuticle). (**F**) Maximum intensity projections of dissected brains of *brp-T2A-QF2^w^* > *QUAS-CD8:GFP* females showing high levels of expression in the entire brain, including antennal lobe glomeruli and higher brain centers. Endogenous GFP projections are separated into anterior (slices 1-22) and posterior (slices 23-50) views (**G**) Maximum intensity projections of dissected ventral nerve cords (VNC) of *brp-T2A-QF2^w^* > *QUAS-CD8:GFP* females showing high levels of expression in all neuromeres. Endogenous GFP projections are separated into anterior (slices 1-12) and posterior (slices 13-49) views. nc82 and merged channels in **(F)** and **(G)** are projections of full z stacks. Scale bars = 100 μm.

Advancements in genetic engineering have led to development of new tools and technologies, now routinely used in insect model and non-model organisms. Among those tools are CRISPR/Cas9^24^, the Q-system of binary expression^25–27^, and Homology Assisted CRISPR Knock-in (HACK)^28^. These tools have enabled the creation of *Drosophila* and mosquito lines with new transgenes and targeted mutations (both knock-in and knock-out), shedding light on the neuronal bases of insect chemosensory-guided behaviors^13,16,23,28,29^. Yet, genetic reagents that target individual mosquito chemoreceptor genes provide only a glimpse of the complexity of the mosquito chemosensory system. As such, the position, number, and identity of the complement of neurons within the chemosensory appendages of a mosquito are unknown. To address this limitation, a pan-neuronal line of *Aedes aegypti* mosquitoes was generated, allowing direct genetic access to all neurons in this species^30^. A similar pan-neuronal genetic reagent in *Anopheles* mosquitoes does not exist. The creation of genetic reagents in *Anopheles* mosquitoes, even in comparison to other non-model organisms like *Ae. aegypti* mosquitoes, is challenging and limited primarily due to low genetic transformation efficiency and survival. Nonetheless, a pan-neuronal *Anopheles* line could provide a roadmap to the number and spatial distribution of neurons within different sensory appendages. Additionally, having access to all neurons would allow the identification of new chemosensory gene targets and global functional analysis of chemosensory neuron responses to important odors, such as attractants and repellents.

Here, using the Q-system and HACK approach, we generate a pan-neuronal driver line for *Anopheles coluzzii* mosquitoes. We targeted the *bruchpilot* (*brp*) gene, which is involved in the structural integrity of neuronal presynaptic active zones. By inserting a *T2A-QF2^w^* cassette before the stop codon of *brp*, we captured the expression pattern of Brp while maintaining its functional expression. This *brp-T2A-QF2^w^* pan-neuronal line enables genetic access to all neurons in the *Anopheles* mosquito. Utilizing a membrane-targeted GFP reporter line, we visualize, spatially map, and quantify neurons in different mosquito appendages. We focus predominantly on female chemosensory appendages relevant for host-searching and biting. By comparing Orco and pan-neuronal populations, we quantify the likely full complement of IR neurons in the antenna. This study presents a comprehensive investigation of the neurons contained within the sensory appendages of the malaria mosquito.

## Results

Mosquitoes have several sensory appendages important for sensing their environment and executing behaviors (**Fig. 1A**). The primary chemosensory appendages on a mosquito are the antenna, maxillary palp, and labella on the proboscis (**Fig. 1B**). The wing margin (**Fig. 1B**) may also contain chemosensory neurons^31^. Information about the full complement of neurons in an appendage could suggest the complexity or range of sensory responses by that appendage. For example, the total number of neurons in the mosquito antenna could indicate a maximum number of olfactory neurons in this tissue. However, the number of neurons that innervate *Anopheles* mosquito appendages is unknown. Neuronal stains could be used to identify neurons in an appendage, but this approach is time consuming, difficult to reproduce across many individuals, challenging to quantify, and could miss many neurons. The optimal approach would be to genetically label all neurons in the *Anopheles* mosquito by using a pan-neuronal driver line. When paired with a fluorescent reporter, such as GFP, this would robustly and reproducibly label all appendage neurons. We utilized such a genetic approach in this work to generate a comprehensive guide of neurons in an *Anopheles* mosquito’s appendages.

### Generation and validation of pan-neuronal driver line

To generate a pan-neuronal driver line in *An. coluzzii* mosquitoes, we used the Q-system of binary expression^27,32^ paired with the Homology Assisted CRISPR Knock-in (HACK) method^28^. We targeted the broadly expressed neural gene, *bruchpilot* (*brp*), which is involved in the structural integrity and function of neuronal synapses^33–35^. Antibodies to Bruchpilot are often used to label all neuropil in *Drosophila* and mosquito brains^32,36^. Recently, the *brp* gene was targeted to generate a pan-neuronal driver line in *Ae. aegypti* mosquitoes^30^, while the HACK approach was utilized to generate olfactory co-receptor knock-in lines in *Drosophila melanogaster*^29^. We targeted the last coding exon using two guide RNAs (gRNAs) to insert a *T2A-QF2^w^* cassette with a *3xP-mCherry* fluorescence eye selection marker before the stop codon of the *brp* gene (**Fig. 1C**). The T2A is a self-cleaving peptide that induces ribosomal skipping, thus allowing two proteins to be produced from the same transcript; in this case a full length Brp protein (with a small T2A tag) that localizes to synapses and a functional QF2^w^ transcription factor that enters the nucleus. By using this targeted knock-in approach, we captured the endogenous expression pattern of the *brp* gene without disrupting its normal function. The successful insertion of the HACK construct into the *An. coluzzii* genome was confirmed by PCR amplifying fragments spanning the *brp* gene with and without the knock-in constructs in wildtype and *brp-T2A-QF2^w^* individuals (**Suppl. Fig 1A-B**).

The *brp-T2A-QF2^w^* individuals were crossed to the *QUAS-CD8:GFP* reporter line^32^ to validate the pan-neuronal expression of the knock-in driver line. Endogenous GFP expression in the resulting progeny (*brp-T2A-QF2^w^* > *QUAS-CD8:GFP*) was evident through the cuticle and easily observed in the brains and ventral nerve cords (VNCs) of larvae (**Fig. 1D**) and various body regions of the adults (**Fig. 1E**). We next used immunohistochemistry to examine brains (**Fig. 1F**) and VNCs (**Fig. 1G**) of brp>CD8:GFP adult females stained with antibodies against Brp (nc82) to assess the overlap between the *brp-T2A-QF2^w^* line and endogenous Brp expression. We observed broad GFP expression in the entire brain and VNC of the brp>CD8:GFP females, with the signal co-localized with anti-Brp staining. Major regions and structures of the brain were well labeled and distinguishable in the anterior (e.g., antennal lobes and glomeruli, mushroom bodies) and posterior (e.g., central complex, fan-shaped body, subesophageal zone) views (**Fig. 1F**). A strong GFP signal was also observed in individual neuromeres of VNCs, which contain neurons relaying information to and from the brain, as well as motor and sensory neurons projecting to the rest of the body (**Fig. 1G**). This broad brain and VNC labeling was consistent among individuals, and not observed in mosquitoes containing the *QUAS-CD8:GFP* reporter alone or in the *orco-T2A-QF2* > *QUAS-CD8:GFP* line (**Suppl. Fig 1C-F**). These data suggest that the *brp-T2A-QF2^w^* driver can serve as a robust pan-neuronal marker in *Anopheles* mosquitoes.

### Quantification and characterization of *brp-T2A-QF2^w^* expression in the sensory appendages of the mosquito head

We next characterized the labeling of the adult brp>CD8:GFP female peripheral nervous system, starting with the sensory appendages of the head (**Fig. 1A** and **1B**). We observed strong labeling of neuron cell bodies, nerve bundles, and dendritic projections into sensilla in antennae (**Fig. 2A**), maxillary palps (**Fig. 2B**), and labella (**Fig. 2C**). We were also able to visualize the neurons of the labrum (the stylet used for blood feeding) extended out of the labium (the sheet housing the bundle of stylets) of the mouthpart (**Fig. 2D**). These images allowed us to generate a comprehensive accounting of neurons in *Anopheles* chemosensory appendages.

**Figure 2.**
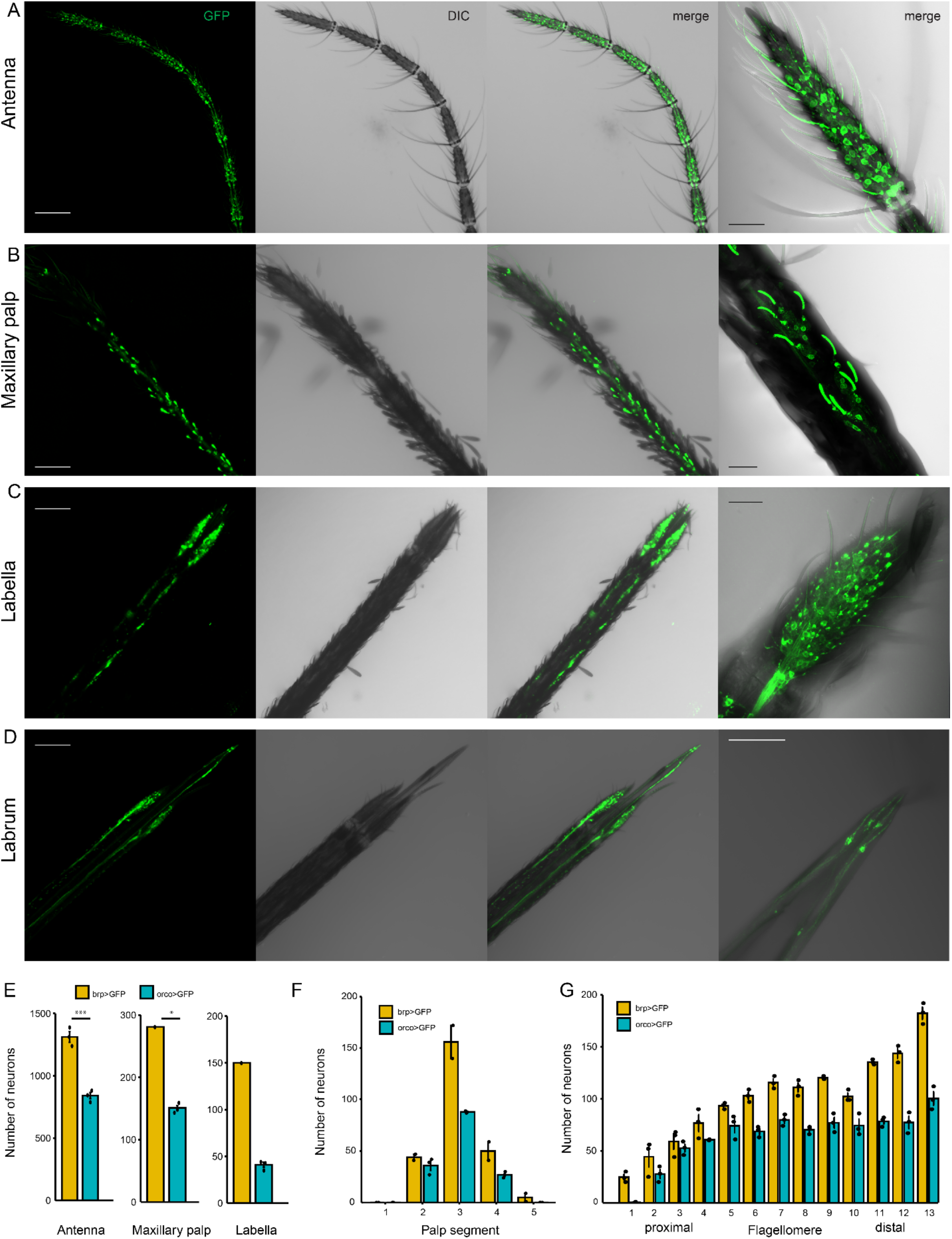
GFP expression in head sensory appendages of *brp-T2A-QF2^w^* > *QUAS-CD8:GFP* female *An. coluzzii*. Maximum intensity projections of z stacks from (**A**) antennae, **(B**) maxillary palps, (**C**) proboscis with labella, and (**D**) labrum (blood-feeding stylet). Images on far right: higher magnification of (A) single antennal segment, (B) section of the maxillary palp, (C) tip of the proboscis (labella), and (D) tip of labrum. Images of individual appendages are obtained from different female mosquitoes. (**E**) Mean number of neurons in a whole antenna, whole maxillary palp, and labellar lobe. **(F)** Mean number of neurons in individual segments of the maxillary palp. **(G)** Mean number of neurons in individual flagellomere of the antenna. **Genotypes of female mosquitoes:** brp>GFP: *brp-T2A-QF2^w^* > *QUAS-CD8:GFP;* orco>GFP: *Orco-T2A-QF2* > *QUAS-CD8:GFP*. Scale bars in A-D are 100 μm (first 3 panels) and 25 μm (far right panels). Statistical comparison in (E) based on independent samples t-test (α= 0.05) *** p< 0.001, * p< 0.5.

While the endogenous GFP signal in the appendages of brp>CD8:GFP individuals was sufficient for visualization of neurons, the quantification of the total number of neurons in those appendages was challenging due to image contrast differences among very brightly and very dimly labeled cells, and the overlap of densely packed neurons in large datasets. To overcome these challenges and to count neurons more efficiently and consistently, we developed a semi-automatic pipeline for image pre- and post-processing to optimize 3-dimensional (3D) neuronal counting (**Suppl. Fig. 2**). The resulting processed images had enhanced contrast and masked any unwanted features that could interfere with counting. The enhancement of the GFP signal in images processed with our pipeline enabled us to detect and count ~42% more neurons in individual antennal segments (n=6) compared to manual counts of unprocessed images (data not shown).

Using our pipeline, we processed images of individual segments of antennae, maxillary palps, and labellar lobes of *brp>CD8:GFP* and *orco>CD8:GFP* female *An. coluzzii* and counted the neurons in those appendages. A single antenna of *brp>CD8:GFP* female mosquito contained an average of 1311 ± 43 (mean ± SE) neurons, 841 ± 23 (mean ± SE) (64%) of which were Orco-positive (Orco+) (**Fig. 2E**). Interestingly, the number of neurons was not distributed equally across the individual segments of the antenna (**Fig. 2G**). Specifically, the total number of neurons increased from the proximal (closest to the head) to distal (the tip) segments of the antenna. This increase in number of neurons was not due to Orco+ neurons, since their numbers stayed mostly unchanged throughout the entire antenna, with the exception of the most distal and most proximal segments. While the identity of the Orco-negative antennal neurons is unknown, they most likely belong to neurons that express the IR olfactory receptor family, as reported by prior RNAseq studies^37^ and recent *in situ* analysis^38^. Additionally, flagellomere segment 1 (the most proximal segment) contains only Orco-negative neurons, which likely include the ~22 Ir93a+ receptor neurons^19^.

A single female maxillary palp contained an average of 255 ± 26 (mean ± SE) neurons, and a single female labellar lobe contained 150 neurons. Maxillary palps and labella of *brp>CD8:GFP* females also contain large populations of Orco-negative neurons (**Fig. 2E**). These Orco-negative neurons account for approximately 41% and 73% of neurons in the maxillary palps and labella, respectively. We also observed neurons in the terminal (most distal) segment of the maxillary palps in the *brp>CD8:GFP* females, but not in *orco>CD8:GFP* females (**Fig. 2F**). The large number of Orco-negative neurons detected in the maxillary palp and labella most likely belong to neurons expressing IR and GR gene families^37,39^.

### Quantification and characterization of *brp-T2A-QF2^w^* expression in the sensory appendages of the mosquito body

We also investigated the GFP expression in female sensory appendages that have not been well characterized (**Fig 1A**), including legs (**Fig. 3A-C**), wings, and the ovipositor. We found that prothoracic (front), mesothoracic (middle) and metathoracic (hind) legs of *brp>CD8:GFP* females contain a large number of neurons (**Fig. 3 A-C**), but only in several of the tarsal segments (tarsomeres). We focused our investigation on terminal (most distal) tarsomere 5, as this segment likely provides important tactile and chemosensory information to the female during contact interactions with plant and animal hosts. Tarsomere 5 of each individual front, middle and hind leg contained on average ~42 neurons (**Fig 3E**), none of which were Orco+ (**Fig 3D**). We also observed labeling of nerve fibers and dendritic projections of some tarsal neurons extending into sensilla. While we did not quantify the number of neurons in the wings or the ovipositor, we observed innervation along the wing margins and across most of the ovipositor surface in *brp>CD8:GFP* females (data not shown). We did not find Orco+ neurons on wings or the ovipositor. Taken together, our results identified the number of neurons in chemosensory appendages with implications on the chemosensory receptor families they may express.

**Figure 3.**
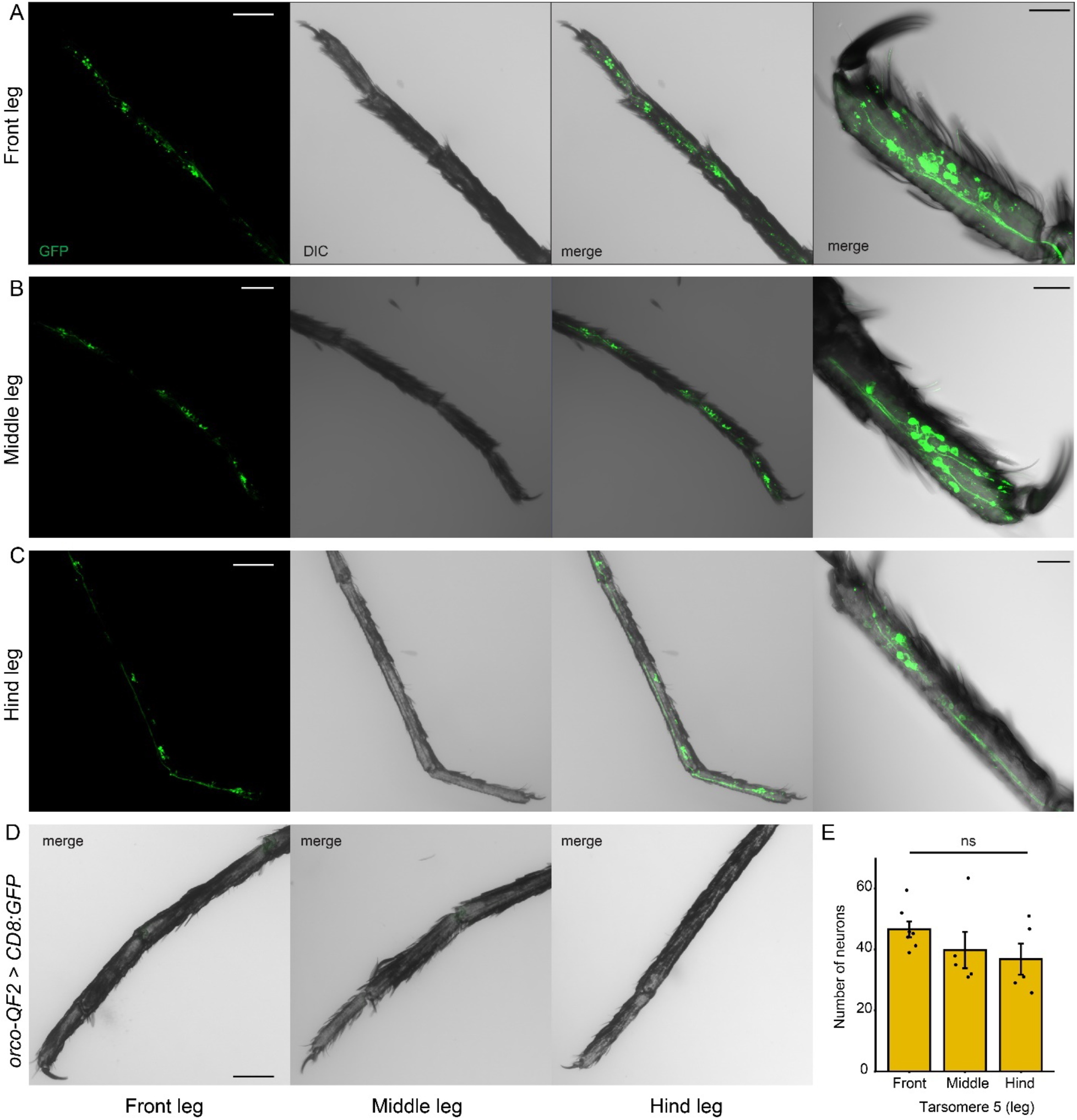
GFP expression in body sensory appendages of *brp-T2A-QF2^w^* > *QUAS-CD8:GFP* female *An. coluzzii*. Maximum intensity projections of z stacks from (**A**) front (prothorasic), **(B**) middle (mesothorasic), (**C**) hind (metathorasic) terminal tarsal leg segments of *brp-T2A-QF2^w^* > *QUAS-CD8:GFP* female mosquitoes. Far right in (A-C): higher magnification images of 5^th^ tarsal segment only. **(D)** Maximum intensity projections of z stacks from front, middle, and hind terminal tarsal leg segments of *orco-T2A-QF2* > *QUAS-CD8:GFP* female mosquitoes. (E) Mean number of neurons in 5^th^ tarsomere of front, middle, and hind legs of *brp-T2A-QF2^w^* > *QUAS-CD8:GFP* female mosquitoes. Statistical comparison based on 1-way ANOVA (α= 0.05). Scale bars: 100 μm (first 3 panels in A-D) and 25 μm (far right panels in A-C).

## Discussion

We developed the first, to our knowledge, pan-neuronal driver line in *An. coluzzii* mosquitoes. By targeting the broadly expressed *bruchpilot* synapse gene, we have enabled genetic access to all neurons in *An. coluzzii* mosquitoes. We further demonstrated the utility of the HACK method, which combines a targeting cassette and gRNAs onto a single plasmid, as an efficient strategy of knocking-in constructs into the *Anopheles* mosquito genome. This approach might also increase knock-in efficiencies in other non-model organisms like *Ae. aegypti*. The pan-neuronal driver line utilizes the Q-system of binary expression and can be crossed to different Q-system compatible reporters for various applications to examine the structure and function of neurons in mosquitoes. Potential applications could include markers for the nucleus, RNAi constructs, calcium indicators, or stochastic labelers.

Prior RNAseq data suggest that female antennae predominantly contain ORs and IRs^37^. With only one GR (Gr1) enriched in female antennae^37^, the vast majority of the Orco-negative neurons we detected most likely express IRs. By comparing the number of neurons in *brp*>GFP and *orco*>GFP *An. coluzzii* female antenna, we can speculate on the distribution and abundance of IR-expressing neurons in the antennae. In contrast to Orco^+^ antennal neurons whose numbers remain relatively constant from segment to segment, the number of the proposed IR^+^ neurons increases from proximal to distal antennal segments. This pattern of IR neuron number and distribution in the *An. coluzzii* antennae suggests that IR-expressing neurons are relatively enriched in distal flagellomeres, supporting fluorescent *in situ* hybridization data examining IR co-receptor expression pattens^38^. We also detected between 1 and 9 brp+ but Orco-negative neurons in proximal segment 1. These are likely the hygro- and thermo-sensory neurons that express Ir93a^19^.

The maxillary palps of *An. coluzzi* females also contain a mixture of Orco+ and Orco-neurons. This sensory appendage contains ~67 capitate peg sensilla, each housing 3 neurons: 1 GR+ and 2 Orco+^37,40–42^. These numbers predict a total of ~200 neurons innervating capitate peg sensilla across the entire maxillary palp. The mean number of neurons in maxillary palps of pan-neuronal females (brp>GFP) exceeded that number. With ~255 neurons total, there are an additional 55 neurons in each female palp that might not innervate capitate peg sensilla. Some of these neurons must be innervating campaniform sensilla and sensilla chaetica, which each contain a single mechanosensory neuron^40^. There is only 1 campaniform sensilla on maxillary palps of *An. gambiae*, but likely numerous sensilla chaetica^40,42^. We are not aware of studies that quantified the number of sensilla chaetica on the maxillary palp of any *Anopheles* species, but the maxillary palps of *Ae. aegypti* contain 14 sensilla chaetica^42^. Since each campaniform sensilla and sensilla chaetica are innervated by a single neuron^40^, and if the number of sensilla chaetica are similar in number between *Aedes* and *Anopheles* mosquitoes, then we predict that at least 15 neurons would be represented by these types of sensilla in each *An. coluzzii* maxillary palp. This prediction suggests ~50 palpal neurons might possibly not be associated with known sensilla. Prior RNAseq analysis suggests that at least some of the neurons on the maxillary palps of *An. gambiae* express receptors belonging to the IR gene family^37^. These IRs might be expressed at low levels in GR+ and Orco+ neurons of the capitate pegs, as was reported for *Ae. aegypti*^43^. Alternatively, IR+ neurons could be among the additional ~50 cells not accounted for by known sensilla. Interestingly, there are no capitate peg sensilla on the most distal segment 5, but this segment does contain 1-9 brp+ neurons. Whether segment 5 neurons are mechanosensory or some are chemosensory remains to be determined.

Labella of *An. coluzzii* mosquitoes contain ~150 brp+ neurons per lobe, comprised of both Orco+ and Orco-neurons. These neurons innervate T1 and T2 sensilla, thought to be gustatory and olfactory, respectively^39,42,44^. There are ~30 T2 sensilla that contain ~60 olfactory neurons (2 neurons/ T2 sensillum) per labellar lobe; by our analyses, only 41-45 of these are Orco+. This count suggests that up to 15 neurons in olfactory T2 sensilla might be expressing another family of olfactory receptors, such as IRs. In addition to ORs and IRs, labella also express many GR receptors^23,39^. These receptors are expressed on dendrites of neurons innervating large T1 gustatory sensilla. There are 15 gustatory sensilla per labellar lobe in *Ae. aegypti*^45^ and *Ae. albopictus* (L. Baik; personal communication), while in *An. quadrimaculatus*, 28 total (14 per lobe) are reported^46^. By examining the labella of *An. coluzzii*, we counted 20-21 long hairs per labellar lobe, tapering towards the tip of the labella, which we presume are T1 sensilla. These sensilla match the locations of T1 sensilla used as stereotypic landmarks for T2 SSR recordings^39^. Assuming the typical 5 neurons per gustatory sensillum (4 GR+ and one mechanosensory), there should be 105 neurons innervating T1 sensilla. Combined with the 60 olfactory sensilla, we can expect ~165 neurons in each labellar lobe. The total number of brp+ neurons (150) is below this predicted number (165), suggesting that there may be fewer than 5 neurons in some T1 sensilla.

Leg tarsi are the first appendages that come into direct contact with a plant or animal host. Since none of the neurons in the tarsi were Orco^+^, any chemosensory neurons in *An. coluzzii* legs are likely IR+ or GR+. The majority of sensory sensilla on the tarsi are predicted to be gustatory. Gustatory sensilla typically contain 4 GR+ chemosensory neurons and 1 mechanosensory neuron. Thus, the 42 neurons we observed on the 5^th^ tarsomeres of front, middle, and hind legs might represent 8 gustatory sensilla with 32 chemosensory neurons that directly report on the taste of host skin, plant, or oviposition surfaces. Characterizing the taste profiles of these chemosensory neurons will be of great interest. Sensilla on leg tarsi may also represent targets for contact repellents such as DEET. In *Ae. aegypti*, the contact repellency of DEET was *orco* independent^13^ and required tarsal contact with a DEET treated surface^47^. Similarly, *An. coluzzii* females were not repelled by volatile DEET^48^, but were repelled by DEET when it contacts their legs^49^. Thus, the 42 tarsal neurons identified here might contain receptor neurons that mediate aversion to DEET or other natural or synthetic repellents.

Our neurogenetic investigation allowed us to identify neurons in less studied and poorly understood sensory appendages of *An. coluzzii* female mosquitoes. While we did not quantify the number of neurons in wings and ovipositors, those sensory organs lacked Orco+ neurons suggesting they express other families of sensory receptors. Wings of *Drosophila melanogaster* contain gustatory receptors which respond to sweet and bitter stimuli and detect pheromones involved in courtship behavior^31,50^. Thus, the neurons in wings and ovipositors identified here for *An. coluzzii* might have sensory functions and play roles in mating or oviposition site selection.

We also noted interindividual variation in the number of neurons across different appendages. Among whole antennae from individual females, the number of neurons differed by 70-150 (*brp > CD8:GFP*) and differed by 24-80 for Orco+ neurons (*orco* > *CD8:GFP*), accounting for up to 11% difference in neuronal counts. In palps and legs, the neuronal count among individuals differed by as much as 20% (50 neurons) and 76% (32 neurons), respectively. All else being equal (genetic background, imaging, image processing and neuron counting), this variation likely reflects biological, rather than technical, differences. Phenotypic variation is necessary for a trait to have the ability to respond to natural selection and environmental changes^51,52^. In the mosquito chemosensory context, with appendages that are predominately required for taste and smell, this difference in number of neurons within individuals also suggests a variation in the number of expressed chemoreceptors (i.e., more neurons to express more chemoreceptors). Consequently, individual females could have different chemosensory sensitivities and behavioral responses in the context of the same chemical cues from attractants and repellents. This interindividual neuronal variation may also influence blood-host selection and preferences. Biological variability in chemosensory neuron numbers may afford individual mosquitoes a range of response properties that in turn benefit the adaptability of mosquito populations.

We successfully utilized the QF2 transcription factor to genetically target broad expression in the central and peripheral nervous system. The viability of the *brp-QF2^w^* line suggests the Q-system can be an appropriate method to target any desired neuronal population in *Anopheles* mosquitoes. In this work, we targeted *T2A-QF2^w^* before the stop codon of *bruchpilot* to allow full-length Bruchpilot to be expressed along with QF2. Similar approaches can be used to maintain the function of other genes while capturing their expression patterns. Alternatively, *T2A-QF2* (or just *QF2*) could be used to target near the start of the gene’s coding region. This would disrupt the function of the gene while still using the Q-system to capture its expression pattern. This would be particularly useful to positively label the cells that contain the mutated gene.

The pan-neuronal line developed here opens up many new avenues of *Anopheles* mosquito research. For example, the pan-neuronal line along with a genetically encoded calcium indicator^53^ enables identification of the neurons in the *Anopheles* mosquito that functionally respond to a variety of sensory stimuli, including various chemosensory attractants and repellents that influence host-seeking behaviors. Conversely, if further combined with an *orco* mutant^14^, it could be used to create a strain of *Anopheles* mosquitoes whose functional olfactory neurons were likely to be IR-expressing. This work represents an important step in the development of genetic tools for characterizing the biology of *Anopheles* mosquitoes.

**Supplementary Figure 1.**
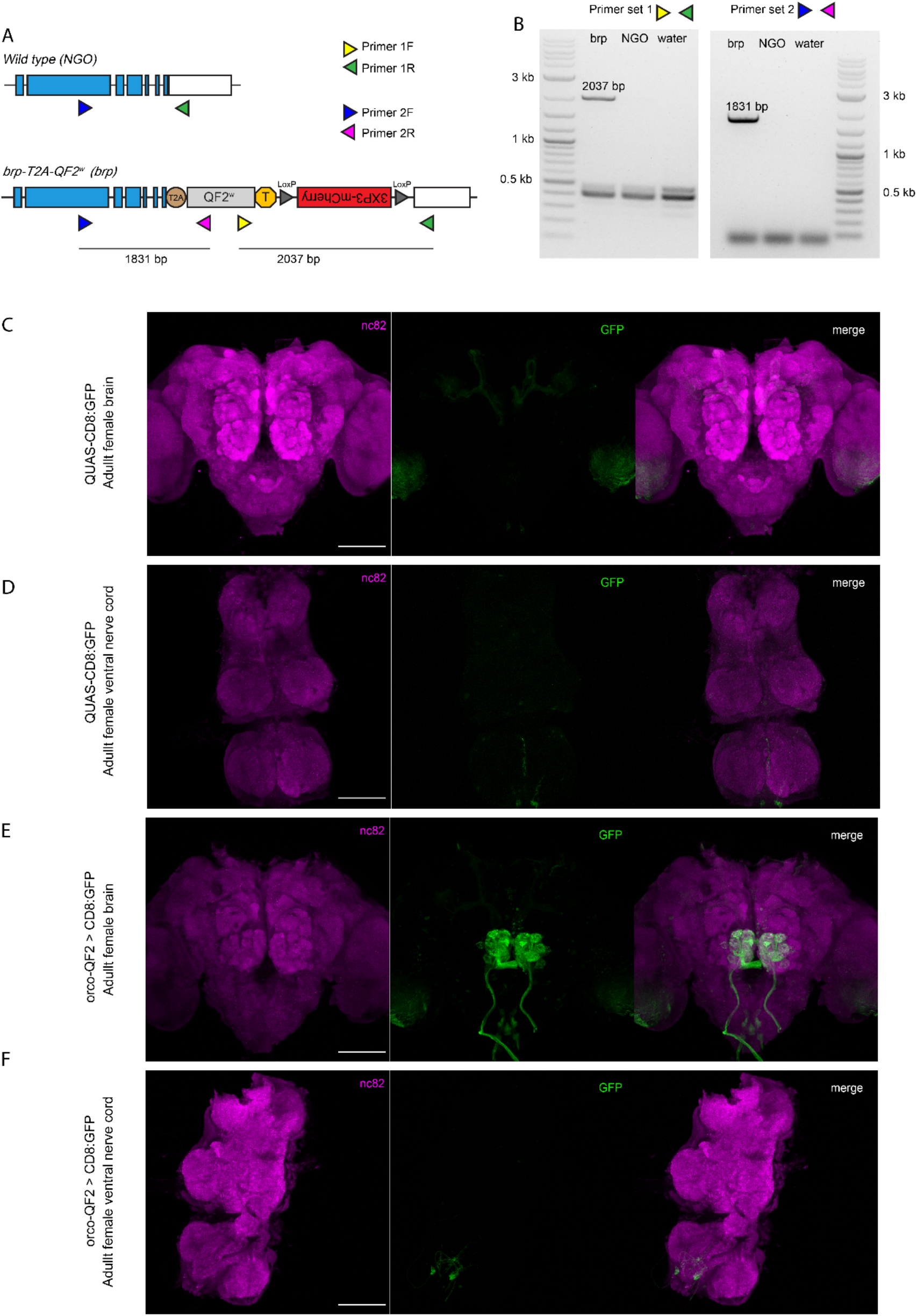
Genotyping and further verification of *brp-T2A-QF2^w^* HACK knock-in driver line. **(A)** Schematic of a section (the last 7 exons) of the wild-type (NGO) and pan-neuronal HACK knock-in (*brp*) genes indicating location of binding sites for genotyping primers. Primers 2F and 1R bind to both the NGO and *brp* DNA. Primer 2F lies outside the genomic region used in the homology arms of the *brp* HACK. Primers 1F and 2R bind only to the *brp* DNA. **(B)** Gel electrophoresis of NGO and *brp* DNA fragments amplified via PCR using 2 sets of genotyping primers. Maximum intensity projections of dissected brains **(C)** and VNCs (**D**) of *QUAS-CD8:GFP* females showing low levels of GFP expression in the mushroom bodies (as previously reported^32^). Maximum intensity projections of dissected brains **(E)** and VNCs (**F**) of *orco-T2A-QF2* > *QUAS-CD8:GFP* females showing high levels of expression in the glomeruli of the antennal lobes. Scale bars = 100 μm.

**Supplementary Figure 2.**
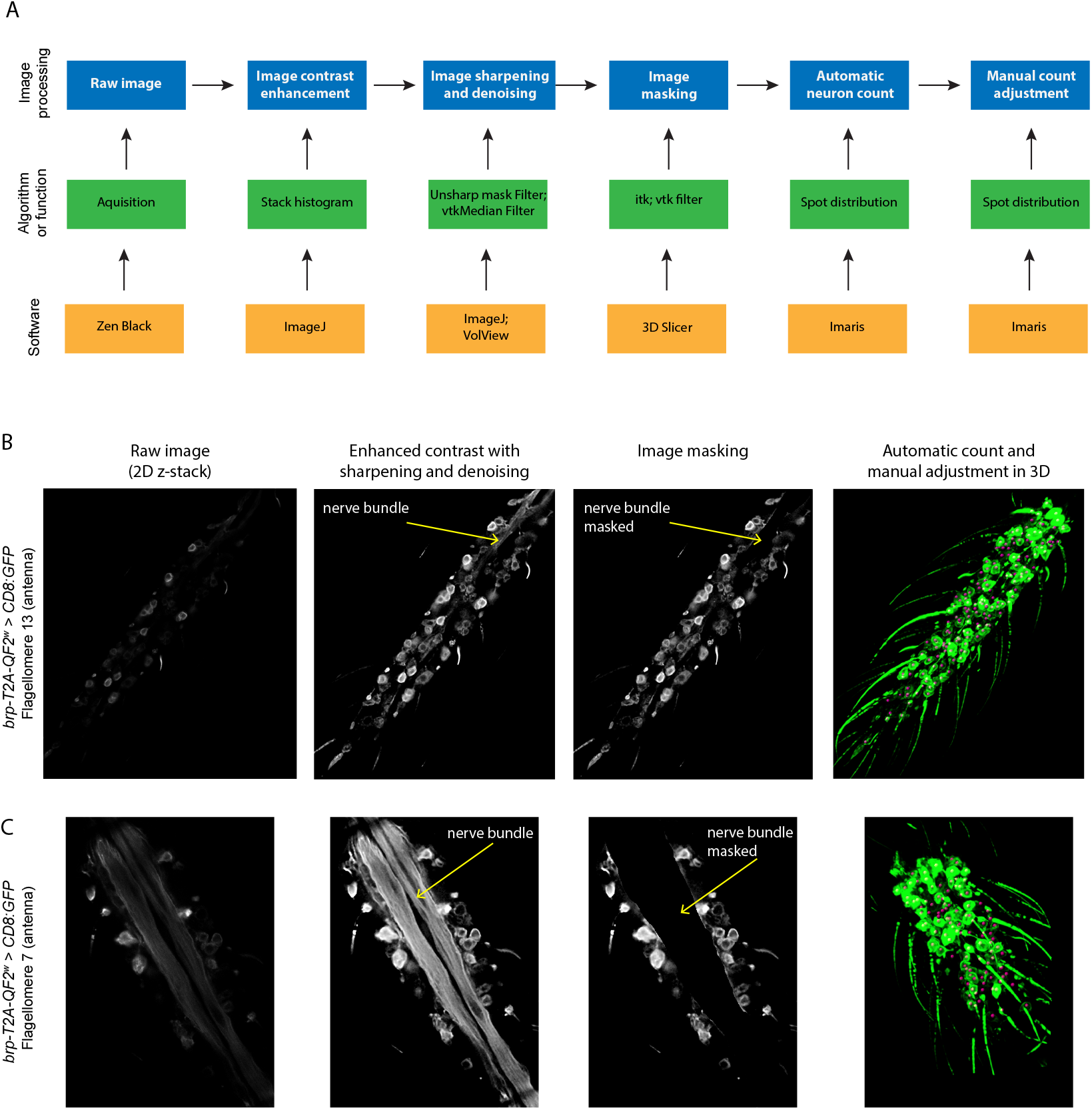
Semi-automatic pipeline for image pre- and post-processing for optimized 3D neuronal counting. **(A)** Schematic of the pipeline workflow indicating the type of image processing (blue), the filters or algorithms used (green), and the software used (yellow). (**B-C**) Examples of 2-D stack confocal images processed via the pipeline. Enhanced contrast with sharpening and denoising, and image masking (masking the nerve bundle) are shown for an image slice of a z-stack of flagellomere 13 **(B)** and flagellomere 7 **(C)** of *brp-T2A-QF2^w^ > QUAS-CD8:GFP* female *An. coluzzii* used as representative images. The pipeline processes the entire z-stack. Automatic count and manual adjustments are shown on a 3D rendered maximum intensity projection with all neurons marked with magenta spheres.

## Acknowledgments

This work was supported by a Natural Science and Engineering Research Council Postdoctoral Fellowship to JKK and by grants from the National Institutes of Health (NIAID R01Al137078), the Department of Defense (W81XWH-17-PRMRP), and the Johns Hopkins Malaria Research Institute (Pilot Fund) to CJP. We thank Barbara Smith (JHU Microscope Facility) and Michele Pucak (JHU Imaging Core Facility) for technical assistance in image acquisition. We thank Chun-Chieh Lin for assistance in cloning and plasmid construction. We thank the Johns Hopkins Malaria Research Institute and Bloomberg Philanthropies for their support.

## Author contributions

JKK and CJP conceived, designed, and interpreted all experiments. JKK and DT designed the HACK construct and performed immunohistochemistry on brains and VNCs. JKK generated and characterized the transgenic line. DP developed the image processing pipeline and processed the imaging data. JKK and DP performed neuronal counts. JKK and CJP wrote the manuscript with inputs from DT and DP.

## Declaration of interests

The authors declare no competing interests

## STAR Methods

### RESOURCE AVAILABILITY

#### Lead contact

Further information and requests for resources and reagents should be directed to and will be fulfilled by the lead contact, Christopher J. Potter.

#### Materials availability

Plasmids generated in this study are available from the lead contact upon request.

#### Data and code availability

Confocal files are available upon reasonable request.

### EXPERIMENTAL MODEL AND SUBJECT DETAILS

#### Mosquito rearing and colony maintenance

Unless otherwise indicated, all wild-type N’Gousso (NGO) and pan-neuronal (brp) *An. coluzzii* mosquitoes were reared at 28±1°C, 80±5 % RH, and 12L:12D light cycle. Eggs were collected on filter paper (Fisherbrand; 9 cm diameter; 09-801B) folded into cones and placed in 3 oz cups filled with reverse osmosis (RO) water. Eggs were hatched into 10 x 12-inch trays (Photoquip Inc. USA) filled with ~ 1L of RO water. Once larvae reached a second instar stage, they were separated and reared at a density of 140-170 individuals/tray until pupation. Larvae were fed daily with ground up TetraMin tropical fish food flakes supplemented with cat food pellets (Cat Chow, Purina) once a week. Collected pupae were allowed to eclose directly into 6 × 6 × 6-inch aluminum cages (BioQuip Products Inc). Adults were supplied with *ad libitum* 10% sucrose solution. Five- to 14-day-old mosquitoes were blood fed on ketamine-anesthetized Swiss Webster mice for 15 minutes or until at least 5-10 females were observed to be fully engorged on blood.

### METHOD DETAILS

#### Plasmid construction and HACK transgenesis

All cloning was performed using In-Fusion Cloning (Clontech #639645). Mosquito genomic DNA was extracted using the DNeasy Blood and Tissue Kit (Qiagen #69506). Cloning steps were confirmed by PCR genotyping (Phusion, NEB) and Sanger sequencing (Genewiz).

##### Construction of Anopheles coluzzii HACK Backbone

The *Anopheles coluzzii* HACK backbone was created using the original *Drosophila melanogaster* HACK construct (Addgene #80274)^28^ with the following modifications. First, the mosquito HACK backbone was designed to have a negative selection marker (GFP) outside of the knock-in homology arms (see **Fig. 1C**), in addition to the positive selection marker (mCherry) within the sequence to be knocked into the mosquito genome. The negative selection marker is used to ensure that the entire plasmid has not been erroneously inserted at an off-target location in the genome. All correctly HACKed mosquitoes should be mCherry+ and GFP-. The *3XP3-GFP-SV40* negative selection marker cassette was constructed from fragments PCR amplified from *pQUAST-mCD8::GFP* (Addgene #24351) and *pXL-BACII-ECFP-15xQUAS-TATA-GCaMP6f-SV40*^48^. In addition, each of the two separate *Drosophila U6* promoters (*U6-1* and *U6-3*) was replaced with a synthetic gBlock (Integrated DNA Technology) for the *Anopheles gambiae U6* short promoter based on the *AgU6* AnGam-2 sequence from Konet et al.^54^ This *Anopheles U6* short promoter was used in tandem to drive expression from each of the two different gRNAs (*AgU6:gRNA1, AgU6:gRNA2*).

AgaU6+gRNA core (gBlock) (*U6* promoter is italicized; *BbsI* flanked spacer sequence is lowercase; gRNA scaffold is underlined):

CTAGTGATCTGAATTAGATCT*ACGGCTGCGTGTGGCTTCTAACGTTATCCATCGCTAGAAGT GAAACGAGCGTGCGTAGGTATATATATGAAATGGAGTTGCTCTCTGCTT*ggggtcttcgagaagacct <u>GTTTTAGAGCTAGAAATAGCAAGTTAAAATAAGGCTAGTCCGTTATCAACTTGAAAAA GTGGCACCGAGTCGGTGC</u>TTTTTTGCCTACCTGGAGCCTGAGAGTTGTTCAATTAATTAA TTCTGACGTAAG

##### Construction of Acol QF2^brp-HACK^ and QF2w^brp-HACK^ Plasmids

We used the HACK approach to target the gene *bruchpilot* (*brp*) to produce a pan-neuronal *Anopheles coluzzii* knock-in line. The homology arms and gRNAs were designed based on the reference sequence for this gene region from Vectorbase (https://vectorbase.org/; ACON010286), which we checked with sequencing against our lab wildtype NGO strain to adjust for potential SNPs. We selected gRNAs by analyzing the region around the stop codon with https://flycrispr.org/^55^ using the following criteria: one gRNA targeting upstream of the stop codon, the second targeting downstream of the stop codon, the two gRNAs being <100 bp apart, and minimizing predicted off-target cleavage sites. The following gRNAs were selected (PAM sequence in parentheses):

gRNA1 AAGCTCTTGAGGAAACCTGC(TGG)
gRNA2 TTTAAGTAAGACCCAGTTAT(TGG)

Several synonymous nucleotide substitutions were made in the homology arms to prevent these gRNAs from targeting the donor sequence. Homology arms were PCR amplified from genomic DNA, while the gRNAs were amplified from the HACK backbone, with the primers themselves adding in the gRNA sequences in place of the spacer sequence in the backbone. The following primers were used for PCR amplification and In-Fusion cloning (bold indicates the In-Fusion 15 base pair overhangs, while underline indicates the synonymous base pair substitutions):

brp_gRNA_FOR: **GTTGCTCTCTGCTTG**AAGCTCTTGAGGAAACCTGCGTTTTAGAGCTAGAAATAGCAAG TTA
brp_gRNA_REV: **TTCTAGCTCTAAAAC**ATAACTGGGTCTTACTTAAACAAGCAGAGAGCAACTCC
brp_5HA_FOR: **ATCGTCGAGTGGTAC**GTGCATTGATTTTGGGTTAGTAATTGCTTGTTTTCTTC
brp_5HA_REV: **GCCCTCACGCGTTAC**GAAGAAGCTCTTGAG<u>A</u>AA<u>G</u>CC<u>G</u>GCTGGTC
brp_3HA_FOR: **AATTAGATCTCTCGA**TGCGACTAGAAACAAAAAACAACTACAAT
brp_3HA_REV: **ACGCAGCCGTCTCGA**GGTACAAATAGCGATTACACACTTGCC

The 5’ homology arm was 1040 base pairs, while the 3’ homology arm was 1075 base pairs. The mosquito HACK backbone was digested with SnaBI to clone in the 5’ homology arm, XhoI to clone in the 3’ homology arm, and BbsI to clone in the gRNAs.

The original HACK backbone (both for flies and mosquitoes) uses the transcriptional activator QF2. Given the expected broad expression of a pan-neuronal driver, we decided to use *QF2^w^* which had also been used for a pan-neuronal driver line in *Ae. aegypti*^30^. We therefore replaced the *QF2* sequence with *QF2^w^* (weak variant) by digesting *Acol QF2^brp-HACK^* with SfiI and AflII, and subcloning in the PCR amplified *QF2^w^* sequence from *pattB-nsyb-QF2^w^* (Addgene #46116).

#### Knock-in Line Establishment and verification

##### Microinjections and Knock-in verification

Plasmids for injections were prepared using ZymoPURE II Plasmid Maxiprep Kit (Zymo Research, USA) and eluted in ultrapure water. To create a microinjection mix*, brp-T2A-QF2^w^* plasmid was mixed with a helper plasmid containing *Cas9* (PX165, pSpCas9, Addgene #48137) and 10x microinjection buffer (50mM KCl, 1mM NaPO4, pH 6.8). A total of 500 ng/μL of DNA (1:1 ratio of plasmids, 250 ng/μL each) was used in microinjection buffer (at final concentration of 1x). Prior to injections, the plasmid mix was passed through 0.22 filter (Ultrafree MC Centrifugal Filter, Merck Millipore Ltd; UFC30GV0S).

Microinjections were performed using an adapted method for mosquito germline transformation^56–58^. Briefly, three to four days following blood-feeding, *An. coluzzii* were offered a lid of a 50 mL conical tube filled with a thin layer of RO water, lined with filter paper (Fisherbrand; 3.5 cm diameter; 09-801-BB) and placed in the dark for 15 minutes to allow for oviposition. Using a fine brush (Winstonia Kolinsky Sable Nail-Art Detail Brush #0000), freshly laid eggs were lined up lengthwise against the edge of a cut nitrocellulose membrane (BrightStar Plus; Invitrogen: AM10100) placed on a glass microscope slide with their posterior poles facing up. A piece of cut and moistened filter paper (Fisher Scientific; 09-801B) was laid on top of the membrane to keep the eggs hydrated. Quartz filaments (Sutter Instruments; QF100-70-10) were pulled into injection needles using a P-2000 micropipette puller (Sutter Instruments) with the following paraments: Heat = 700; Vel = 60; Del = 145; Pul = 175. Eggs were injected under an Olympus SZX16 microscope with Olympus SDF PLAPO 1.6xPF lens using FemtoJet and PatchManNP2 (Ependorf North America) micromanipulator. Injection parameters used were: 300-1200 hPa injection pressure (adjusted as needed depending on the opening of the needle), 100-500 hPa back pressure, and 0.1-0.2 s injection time. Injected eggs were kept hydrated and left undisturbed until they were fully melanized, at which point they were transferred to a hatching platform (cut and inverted 25 ml polystyrene reagent reservoir wrapped in paper towel) and placed in a 12.5 × 8.5 × 7 cm plastic container with ~ 120 mL of RO water. A total of 380 eggs were injected and 33 of those hatched.

Surviving adults were separated by sex at pupal stage and crossed *en masse* to NGO individuals of the opposite sex. Crosses were blood-fed 4 times with all progeny screened using an Olympus SZX7 epifluorescence microscope equipped with RFP (for mCherry eye marker) and GFP filters. Animals were illuminated with an X-Cite Series 120Q light source. Images were acquired using a QImaging QIClick Cooled digital CCD camera and Q-Capture Pro 7 software. In total, 15 G_1_ larvae (out of 1364 screened) were positive for the red eye marker (mCherry) and negative for green (GFP) eye marker, indicating that the construct was integrated at the targeted location. The individual larvae that survived to adult stage were outcrossed to NGO individuals of the opposite sex in 1:3-5 ratio. A single cross from one G_1_ female produced viable eggs, making this individual a single founder of the *brp-T2A-QF2^w^* pan-neuronal line. The knock-in was confirmed by PCR genotyping (Phusion, NEB) using primers that bind outside and inside the brp knock-in region (**Supplementary Fig. 1A-B**). The following primers were used to genotype the mosquitoes:

Primer 1F CTCTCGATGCTATCACTCAGACCAA
Primer 1R TTCTCAATTGAAGCTAGCAGCAACC
Primer 2F GCTGAAACAAATGCTTCAGGAAACG
Primer 2R TGTATTCCGTCGCATTTCTCTC

The amplified fragments were 2037 base pairs and 1831 base pairs from primer set 1 and primer set 2, respectively.

#### Reporter expression

Expression was examined by crossing to a *QUAS-mCD8::GFP* reporter line previously established by our lab^32^.

##### Immunohistochemistry

Brain and ventral nerve cord (VNC) staining was carried out as previously described^32,59^. Briefly, bodies of 9-25 dpe female mosquitoes were fixed in 4% paraformaldehyde in 0.1 M Millonig’s phosphate buffered solution (pH 7.4) (Electron Microscopy Sciences: 11582-05) for 3-4 hours at 4°C. Brains and VNCs were dissected out in 1xPBS and washed in PBT (1x PBS containing 0.3% Triton X-100 (Electron Microscopy Sciences: 22140)) for 1 h (3 times, 15-20 min each) at room temperature (RT). The tissues were then permeabilized with a blocking solution (1x PBS containing 4% Triton X-100 and 2% normal goat serum (NGS)) overnight at 4°C. The following day, brains and VNCs were washed for 1h in PBT (3 times, 15-20 min each) at RT and incubated in PBT with 2% NGS with primary antibodies for 3 nights at 4°C. The primary antibodies used were rat anti-CD8 (Invitrogen #MCD0800, 1:100) and mouse anti-nc82 (DSHB, AB_2314866, 1:50). After incubation with primary antibodies, tissues were washed for 1h in PBT (3 times, 15-20 min each) at RT and incubated in PBT with 2% NGS with secondary antibodies for 3 nights at 4°C. The secondary antibodies used were Cy3 goat-anti rat (Jackson ImmunoResearch #112-165-167, 1:200) and Alexa-647 goat anti-mouse (Life Technologies Z25008, 1:200). After incubation in secondary antibodies, brains and VNCs were washed for 1 h in PBT, placed in a mounting solution (*SlowFade* Diamond Antifade Mountant; Life Technologies Corporation, S36972) overnight at 4°C, and mounted on microscope slides (Thermo Scientific, 3050-002) the following day. Endogenous CD8:GFP expression in the mounted tissues was visualized using laser confocal microscopy.

##### Confocal imaging

Female mosquitoes (≥ 7 dpe) were briefly chilled on ice. Individual sensory appendages were removed with a pair of forceps or a fine microblade and mounted on microscope slides in a drop of a mounting solution (*SlowFade* Diamond Antifade Mountant; Life Technologies Corporation, S36972). Legs were mounted directly after being detached from the bodies. To minimize air bubbles, antennae, palps, and labella were incubated in a fixative solution (4% paraformaldehyde in 0.1 M Millonig’s phosphate buffered solution) for 5 minutes prior to being mounted on slides. Imaging of sensory appendages was done within 2 hours of detaching from the body to capture the endogenous GFP signal.

Images of brains, VNCs, and sensory appendages were obtained using a Zeiss LSM 700 confocal microscope equipped with Fluar 10×/0.50 air M27, LCI Plan-Neofluar 25×/0.8 water Korr DIC M27, and Plan-Apochromat 40×/1.3 Oil DIC M27 objectives. Images were acquired at 1024 × 1024-pixel resolution. Images of brains and VNCs were captured with 2.45 μm z-steps using 25x objective. For sensory appendages imaged with 10x and 40x objectives, the z-steps were 2.84 μm and 0.42 μm, respectively. Maximum intensity projections of full z-stacks or partial z-stacks were generated using Fiji/ImageJ.

##### Semi-automatic pipeline for 3-D neuron quantification in mosquito appendages

The acquired z-stack raw images for each appendage were processed through a modified image analysis framework^60^ in Fiji/ImageJ(v.1.53t), VolView (v.3.4), 3D Slicer (v. 5.0.3), and Imaris (v. 9.9.1) to enable neuron counting in 3D. First, image contrast enhancement was performed using the stack histogram method in ImageJ on the z-stack, followed by image sharpening using unsharp mark filter (radius = 3 px) in ImageJ, and image denoising using the vtkmedian filter (kernel size = 3) in VolView. Masking of the nerve bundle in the images across the entire z-stacks was performed in 3D Slicer. Finally, the spots distribution algorithm in Imaris was used to automatically detect all 3D objects of ~3.5 μm diameter. Each count was adjusted manually to remove detected objects that were not neurons (e.g., brightly labelled sensilla), and neurons that were missed (e.g., overlapping neurons) were added. The adjustments were made in 3D rendered maximum intensity projections and verified in 2-D planes.

#### Life history and fitness of *brp-T2A-QF2^w^* mosquitoes

Broad expression of exogenous transcription factor proteins such as GAL4 and QF2 can cause toxicity and lead to behavioral defects and lower fitness in transgenic animals^27,61^. While we used QF2^w^, a weaker version of QF2^27^, we noticed some defects in the pan-neuronal *brp-T2A-QF2^w^* individuals after carefully quantifying several life history traits of this line. It is possible that the T2A peptide added to the C-terminus of the Brp protein might also affect its function.

Attempts at making the pan-neuronal line homozygous were not successful, as pan-neuronal male to female crosses produced few, if any, eggs. We thus kept the pan-neuronal line as heterozygotes by crossing the pan-neuronal individuals *en masse* to wild-type counterparts of the opposite sex each generation. Crosses of pan-neuronal males to wild-type females produced a higher number of eggs than crosses of pan-neuronal females to wild-type males. However, the progenies were equally likely to inherit the transgenic copy of the *brp* gene from both males and females, as we did not detect a difference in the proportion of RFP+ larvae between the two pan-neuronal crosses. This trend was consistent over 10 generations. The female-specific defect can be attributed to low host attraction and blood feeding of the heterozygous pan-neuronal females. When kept in small groups with wild-type males, those females were not attracted to a host (anesthetized mouse) and they were thus unable to successfully blood-feed. Only a small proportion of the heterozygous pan-neuronal females successfully blood-fed when kept in large groups of at least 50-100 individuals. Larva to pupa and pupa to adult survival rates of panneuronal individuals were comparable to wild-type controls, with only a small increase in time to pupation.

### QUANTIFICATION AND STATISTICAL ANALYSIS

Data are reported as means ± standard errors of means (SEM). All statistical analysis was carried out in R v. 4.2.1 (R Core Team 2022). All data were checked for normality with QQ plots and Shapiro-Wilk tests. Homogeneity of variance was checked with variance tests (for comparisons between two groups) or with Bartlett tests (for comparisons among more than 2 groups). Number of neurons between genotypes were compared using independent samples t-tests or 1-way ANOVA. For all *orco>CD8:GFP* n=3. For *brp>CD8:GFP* n=3 (antennae), n=2 (palps), and n=1 (labella). For *brp>CD8:GFP* legs (5th tarsomere), n= 7 (front legs), n= 5 (middle and hind, each). For all statistical comparisons, α was set to 0.05.

## Notes

### Competing Interest Statement

The authors have declared no competing interest.

